# Transformation and model choice for RNA-seq co-expression analysis

**DOI:** 10.1101/065607

**Authors:** Andrea Rau, Cathy Maugis-Rabusseau

## Abstract

Although a large number of clustering algorithms have been proposed to identify groups of co-expressed genes from microarray data, the question of if and how such methods may be applied to RNA-seq data remains unaddressed. In this work, we investigate the use of data transformations in conjunction with Gaussian mixture models for RNA-seq co-expression analyses, as well as a penalized model selection criterion to select both an appropriate transformation and number of clusters present in the data. This approach has the advantage of accounting for per-cluster correlation structures among samples, which can be quite strong in RNA-seq data. In addition, it provides a rigorous statistical framework for parameter estimation, an objective assessment of data transformations and number of clusters, and the possibility of performing diagnostic checks on the quality and homogeneity of the identified clusters. We analyze four varied RNA-seq datasets to illustrate the use of transformations and model selection in conjunction with Gaussian mixture models. Finally, we propose an R package `coseq` (**co**-expression of RNA-**seq** data) to facilitate implementation and visualization of the recommended RNA-seq co-expression analyses.

## Introduction

Increasingly complex studies of transcriptome dynamics are now routinely carried out using high-throughput sequencing of reverse-transcribed RNA molecules (i.e., cDNA molecules), called RNA sequencing (RNA-seq). By quantifying and comparing transcriptomes among different types of tissues, developmental stages, or experimental conditions, researchers have gained a deeper understanding of how changes in transcriptional activity reflect specific cell types and contribute to phenotypic differences. Identifying groups of co-expressed genes may help target gene modules that are involved in similar biological processes [1, 2] or that are candidates for co-regulation. Thus, by identifying clusters of co-expressed genes, we aim both to identify co-regulated genes and to characterize potential biological functions for orphan genes (namely, those whose biological function is unknown).

A great deal of clustering algorithms have been proposed for microarray data, raising the question of their applicability to RNA-seq data. In particular, after normalization, background correction, and log_2_-transformation of microarray data, hybridization intensities are typically modeled by Gaussian distributions [3]. RNA-seq data, on the other hand, are made up of read counts [4, 5] or pseudocounts [6, 7] for each biological entity or feature (e.g., a gene) after either alignment to a genome reference sequence or *de novo* assembly. These data are characterized by 1) highly skewed values with a very large dynamic range, often covering several orders of magnitude; 2) positive correlation between feature size (e.g., gene length) and read counts [8]; and 3) variable sequencing depth (i.e., library size) and coverage among experiments [9]. The presence of overdispersion (i.e., variance larger than the mean) among biological replicates for a given feature is also a typical feature of these data, leading to the use of negative binomial models [10, 11] for RNA-seq differential analyses.

Statistically speaking, the goal of clustering approaches is to discover structures (clusters) within data. Many clustering methods exist and roughly fall into two categories: 1) methods based on dissimilarity distances, including tree-based hierarchical clustering [12] as well as methods like the *K*-means algorithm [13]; and 2) model-based methods [14], which consist of defining a clustering model and optimizing the fit between the data and the model. For the latter class of models, each cluster is represented by a distinct parametric distribution, and the entire dataset is thus modeled as a mixture of these distributions; a notable advantage of model-based clustering is that it provides a rigourous framework to assess the appropriate number of clusters and the quality of clusters obtained. Presently, most proposals for clustering RNA-seq data have focused on the question of grouping biological samples rather than features, for example using hierarchical clustering with a modified loglikelihood ratio statistic based on a Poisson loglinear model as a distance measure [15] or the Euclidean distance of samples following a variance-stabilizing transformation [16].

In recent work [17], we proposed the use of Poisson mixture models to cluster RNA-seq expression profiles. This method has the advantage of directly modeling the count nature of RNA-seq data, accounting for variable library sizes among experiments, and providing easily interpretable clusterings based on the profiles of variation around average expression of each gene. However, there are several serious limitations to this approach: 1) the assumption of conditional independence among samples, given the clustering group, is likely to be unrealistic for the vast majority of RNA-seq datasets; 2) per-cluster correlation structures cannot be included in the model; and 3) the Poisson distribution is likely overly restrictive, as it imposes an assumption of equal means and variances. In addition, classical asymptotic model selection criteria, such as the Bayesian Information Criterion (BIC) [18] and Integrated Completed Likelihood (ICL) criterion [19], were observed to have poor behavior for the Poisson mixture model in many cases. As such, Rau et al. [17] proposed the use of a non-asymptotic penalized model selection criterion calibrated by the slope heuristics [20, 21], requiring a collection of mixture models to be fit for a very wide range of cluster numbers *K*; for large *K*, this can imply significant computational time as well as practical difficulties for parameter initalization and estimation. We note that a related approach based on a hybrid-hierarchical clustering of negative binomial mixtures was proposed by Si et al. [22]; as with the work of Rau et al. [17], this method cannot account for correlation structures among samples.

To address the aforementioned limitations of the Poisson mixture model, in this work we investigate appropriate transformations to facilitate the use of Gaussian mixture models for RNA-seq co-expression analysis. This strategy has the notable advantage of enabling the estimation of per-cluster correlation structures, as well as drawing on the extensive theoretical justifications of Gaussian mixture models [14]. We note that Law *et al.* [23] employed a related strategy for the differential analyses of RNA-seq data by transforming data, estimating precision weights for each feature, and using the `limma` empirical Bayes analysis pipeline [24]. The identification of an “appropriate" transformation for RNA-seq co-expression is not necessarily straightforward, and depends strongly on the desired interpretability of the resulting clusters as well as the model assumptions. Several transformations of read counts or pseudocounts have been proposed in the context of exploratory or differential analyses, but most largely seek to render the data homoskedastic or to reduce skewness. In this work, rather than grouping together genes with similar absolute (transformed) read abundances, we propose the use of normalized expression *profiles* for each feature, that is, the proportion of normalized counts observed for a given feature. Due to the compositional nature of these profiles (i.e., the sum for each feature equals 1), an additional transformation is required prior to fitting the Gaussian mixture model, as discussed below.

The remainder of the article is organized as follows. In the Methods section, we introduce some notation, discuss appropriate data transformation for RNA-seq co-expression analyses, and briefly review Gaussian mixture models, including parameter estimation and model selection. In the Results section, we describe several RNA-seq datasets and illustrate co-expression analyses on each using Gaussian mixture models on transformed data using the `coseq` R package. Finally, in the Discussion we provide some concluding remarks and recommendations for RNA-seq co-expression analyses in practice, as well as some opportunities for future work.

## Methods

For the remainder of this work, let *Y*_*ij*_ be a random variable, with corresponding observed value *y*_*ij*_, representing the raw read count (or pseudocount) for biological entity *i* (*i* = 1,…,*n*) of biological sample *j* (*j* = 1,…,*q*). For simplicity, in this work we typically refer to the entities *i* as genes, although the generality of the following discussion holds for other entities of interest (exons, etc). Each sample is typically associated with one or more experimental conditions (e.g., tissue, treatment, time); to reflect this, let 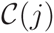 correspond to the experimental group for sample *j*. Finally, let y be the (*n* × *q*) matrix of read counts for all genes and samples, and let y_*i*_ be the *q*-dimensional vector of raw count values across all biological samples for gene *i*. In the following, we use dot notation to indicate summations over a particular index, e.g. *y*_*i*._ = Σ_*j*_ *y*_*ij*_.

### Data transformations for RNA-seq co-expression

A feature common to many RNA-seq data transformations is the incorporation of sample-specific normalization factors, often referred to as *library size* normalization. These normalization factors account for the fact that the number of reads expected to map to a particular gene depends not only on its own expression level, but also 1) on the total number of mapped reads (also referred to as library size) in the sample, and 2) on the overall composition of the RNA population being sampled. Although several library size normalization factors have been proposed since the introduction of RNA-seq, the median ratio [11] and trimmed mean of M-values [TMM; 25] methods have been found to be robust and effective, and are now widely used [26] in the context of differential analysis. Without loss of generality, we note t = (*t*_*j*_) as the scaling normalization factors for raw library sizes calculated using the TMM normalization method; *ℓ*_*j*_ = *y*_*·j*_*t*_*j*_ is then the corresponding normalized library size for sample *j*, and

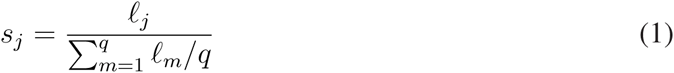

is the associated normalization scaling factor by which raw counts *y*_*ij*_ are divided.

Several data transformations have been suggested for RNA-seq data, most often in the context of exploratory or differential analyses. These include a log_2_ transformation (where a small constant is typically added to read counts to avoid 0’s), a variance-stabilizing transformation [VST; 27, 28, 16], moderated log counts per million [CPM; 23], and a regularized log-transformation [rlog; 11]; see the Supplementary Materials for more details about each. As previously noted, each of these transformations was proposed with the objective of rendering the data homoskedastic (in the case of the VST or regularized log transformations) or to reduce the orders of magnitude spanned by untransformed RNA-seq data. Rather than making use of these transformations, we propose calculating the normalized expression *profiles* for each feature, that is, the proportion of normalized reads observed for gene *i* with respect to the total observed for gene *i* across all samples:

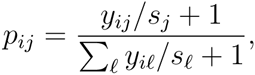

where *s*_*j*_ are the scaling normalization factors for raw counts (see Equation 1). To illustrate the interest of using these normalized expression profiles for co-expression analysis, we plot *y*_*ij*_/*s*_*j*_, log(*y*_*ij*_/*s*_*j*_ + 1), and the normalized expression profiles *p*_*ij*_ in Figure 1 for a subset of genes from the mouse RNA-seq data studied by Fietz et al. [29]. In particular, we consider ten genes that are most representative (as measured by Euclidean distance) of four distinct groups: non-differentially expressed (NDE) genes across all samples (Group 1); genes expressed only in the last experimental condition (samples 11 to 15, Group 2); genes expressed only in the first experimental condition (samples 1 to 5, Group 3); and genes expressed only in the second experiemental condition (samples 6 to 10, Group 4). It may clearly be seen that the large differences in magnitude that are dominant for normalized counts (Figure 1A) are greatly reduced by a log-transformation (Figure 1B), although a certain amount of spread remains between very highly and weakly expressed genes. This spread can be notably reduced by considering the normalized expression profiles *p*_*ij*_ (Figure 1C). This example is thus instructive in illustrating the importance in co-expression analyses of considering a measure that is independent of the absolute expression level of the genes, as is the case for the normalized profiles.

**Figure 1:**
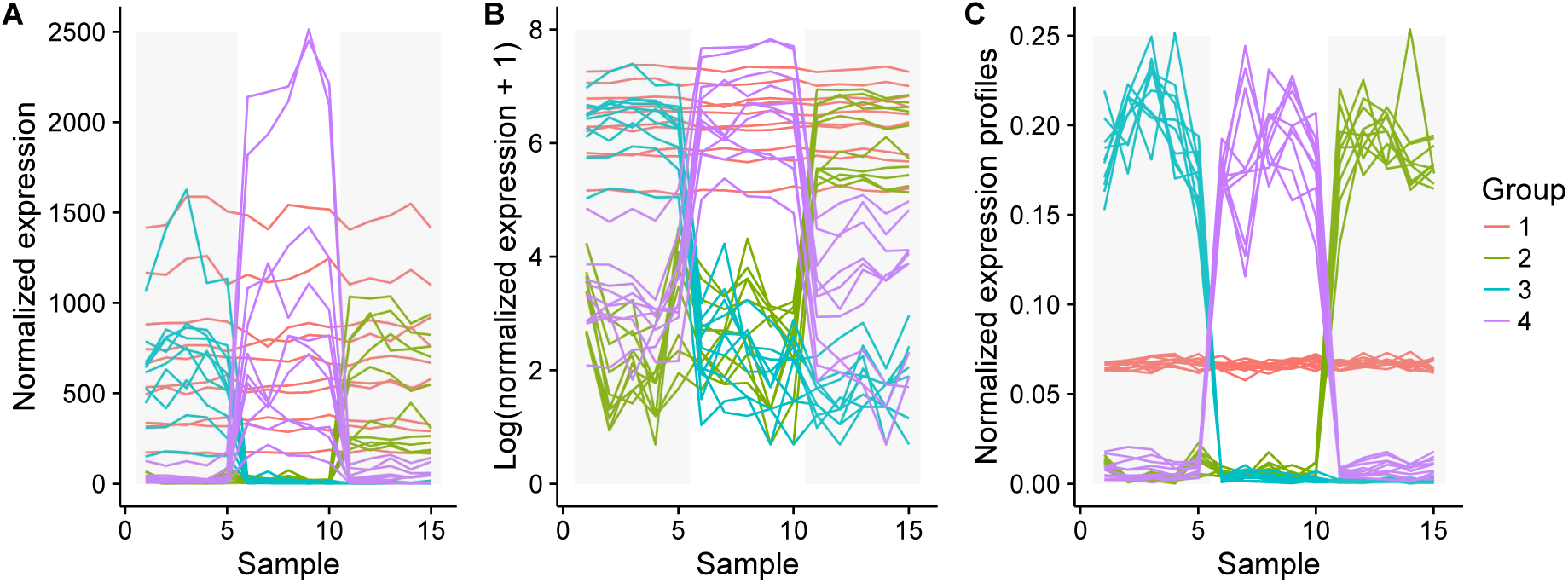
Normalized counts (A), log normalized counts + 1 (B), and normalized expression profiles *p*_*ij*_ (C) for a subset of the Fietz et al. [29] mouse RNA-seq data. The subset of genes include non-differentially expressed (NDE) genes across all samples (Group 1); genes expressed only in the last experimental condition (samples 11 to 15, Group 2); genes expressed only in the first experimental condition (samples 1 to 5, Group 3); and genes expressed only in the second experiemental condition (samples 6 to 10, Group 4). Transparent grey boxes delimit the replicates in each of the three experimental groups.

It is important to note that the profile for gene *i*, **p**_*i*_ = (*p*_*ij*_), represents compositional data [30], as it is a q-tuple of nonnegative numbers whose sum is 1. This means that the vector of values **p**_*i*_ are linearly dependent, which imposes constraints on the covariance matrices Σ_*k*_ that are problematic for the general Gaussian mixture model (and indeed for most standard statistical approaches). For this reason, we consider two separate transformations of the profiles *p*_*ij*_ to break the sum constraint, the logit and the arcsin (also referred to as the arcsin square root, or angular) transformations:

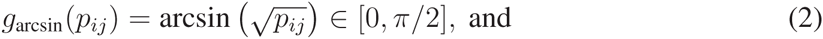

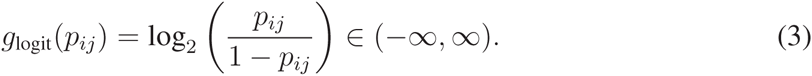

Over a broad range of intermediate values of the proportions, the logit and arcsin transformations are roughly linearly related to one another (see Figure S1A in the Supplementary Materials). However, although both transformations tend to pull out the ends of the distribution of *p*_*ij*_ values, this effect is more marked for the logit transformation, meaning that it is more affected by smaller differences at the ends of the scale (Figure S1B).

### Gaussian mixture models

Model-based clustering consists of assuming that the expression data come from several separately modeled subpopulations, where the full population of genes is a mixture of these subpopulations. Thus, observations are assumed to be a sample from an unknown probability distribution with density *f*, which is estimated by a finite mixture

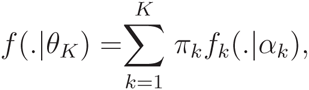

where *θ*_*K*_ = (**π**, *α*_1_,…, *α*_*K*_), and **π** = (*π*_1_,…, *π*_*K*_) are the mixing proportions, with *π*_*k*_ ∈ (0,1) for all *k* and 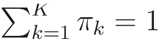. The density *f*_*k*_ (·|*α*_*k*_) of the *k*^th^ subpopulation must be chosen according to the nature of the gene expression measures; in the following, we consider the special case of Gaussian mixture models.

A collection of Gaussian mixture models can be defined as 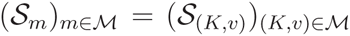, where

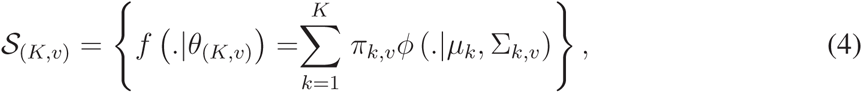

with *ϕ* (*·*|*μ*_*k*_, Σ_*k,v*_) denoting the *q*-dimensional Gaussian density with mean *μ*_*k*_ and covariance matrix Σ_*k,v*_. The index *v* denotes one of the Gaussian mixture shapes obtained by constraining one or more of the parameters in the following decomposition of each mixture component variance matrix:

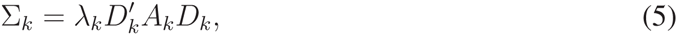

where *λ*_*k*_ = |Σ_*k*_|^1/*q*^, *D*_*k*_ is the eigenvector matrix of Σ_*k*_, and *A*_*k*_ is the diagonal matrix of normalized eigenvalues of Σ_*k*_. Various constraints on these parameters respectively control the volume, orientation, and shape of the *k*^th^ cluster [31]; by additionally allowing the proportions *π*_*k*_ to vary according to cluster or be equal for all clusters, we may define a collection of 28 parsimonious and interpretable mixture models, available in the `Rmixmod` R package [32]. Without loss of generality, for simplicity of notation we will consider here only the most general model form, with variable proportions, volume, orientation, and shape (referred to as the [*p*_*k*_*L*_*k*_*C*_*k*_] in `Rmixmod`); as such, the model collection is defined solely over a range of numbers of clusters, 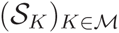.

The parameters of each model 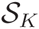 in the collection defined in (4) may be estimated using an expectation-maximization (EM)-type algorithm [33]. After solving the density estimation problem, for each model in the collection *f* is estimated by 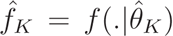, and the data are clustered using the maximum a posteriori (MAP) rule: for each *i* = 1,…, *n* and each *k* = 1,…,*K*,

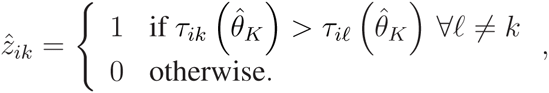

where 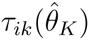 is the conditional probability that observation *i* arises from the *k*^th^ component mixture 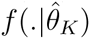.

### Model choice for RNA-seq co-expression

In the mixture model framework, the number of clusters *K* is typically chosen from the model collection using a penalized selection criterion such as the BIC, [18], ICL [19], or a non-asymptotic penalized criterion whose penalty is calibrated using the slope heuristics (SH) principle [34]:

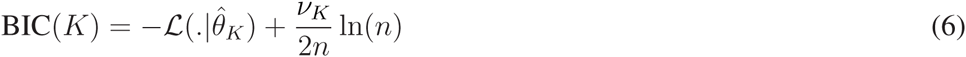

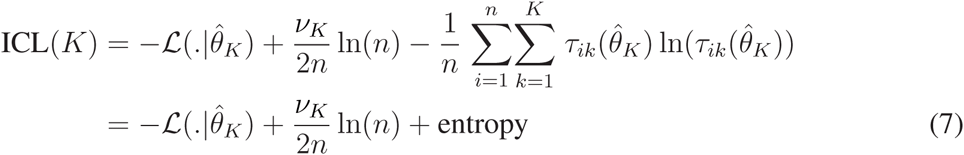

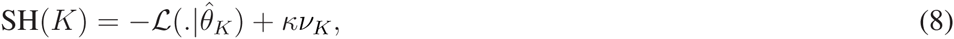

where 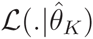 is the loglikelihood evaluated at 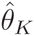 (the maximum likelihood estimate of *θ*_*K*_), *ν*_*K*_ represents the number of free parameters for the mixtures in model 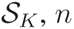 is the number of observations, *κ* is a multiplicative constant that must be calibrated using the `capushe` R package [21], and 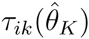 is as defined in the previous section. The selected model 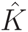 corresponds to the number of clusters *K* that minimizes the chosen criterion among Equations (6–8).

Although rarely done in practice, penalized criteria like the BIC and ICL may also be used to select among different models or transformations, as was suggested in a different context by Thomas et al. [35] and more recently for RNA-seq data by Gallopin [36]. This is of great interest, as it removes the need for an arbitrary choice of data transformation by using the framework of formal model selection. We illustrate this principle for the choice of number of clusters *K* and data transformation; in a more general case, a similar procedure could be used to additionally choose among the different forms of Gaussian mixture models described in Equation (5) or among different parametric forms of models. Let *g*(x) represent an arbitrary monotonic transformation of a dataset x. If the new sample *g*(x) is assumed to arise from an i.i.d. Gaussian mixture density, *f* (.|*θ*_*K*_), then the initial data x is an i.i.d. sample from density *f*_*g*_(.|*θ*_*K*_), which is a transformation of *f* (.|*θ*_*K*_) and thus not necessarily a Gaussian mixture density. If *J*_*g*_ denotes the Jacobian of the transformation *g* and 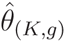 the maximum likelihood estimate obtained for the model with *K* clusters and transformation *g*, we select the pair (*K, g*) leading to the minimum of the corrected BIC or ICL criteria:

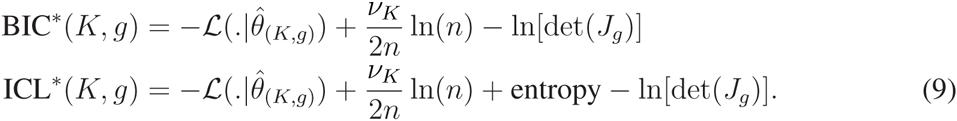

Note that in these expressions, the number of parameters *ν*_*K*_ does not depend on the transformation *g*.

For the purposes of this work, we make use of the corrected ICL criterion defined in Equation (9) to compare between the logit and arcsin transformations in Equations (2) and (3) applied to the expression profiles p = (*p*_*ij*_). In particular, we use the following:

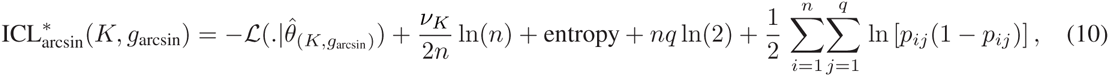

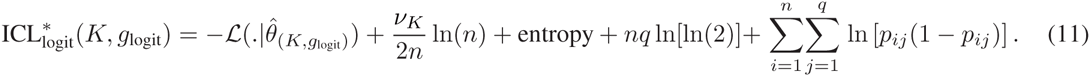

The values of 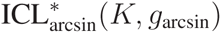 and 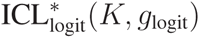 can thus be directly compared to choose between the two transformations.

### coseq R package

To facilitate co-expression analyses of RNA-seq data using Gaussian mixture models and an appropriate data transformation, we have created the R package `coseq` (co-expression of RNA-seq data), freely available on GitHub; in this section, we briefly describe some of the options available in this package.

The package is first installed and loaded via the `devtools package` using the following commands:

~~~
> library(devtools)
> install_github("andreamrau/coseq")
> library(coseq)
~~~

A typical call to `coseq` to fit a Gaussian mixture model on arcsin- or logit-transformed normalized profiles takes the following form:

~~~
> run_arcsin <- coseq(counts, K=2:10, model="Normal", transformation="arcsin")
> run_logit <- coseq(counts, K=2:10, model="Normal", transformation="logit")
~~~

where `counts` represents a (*n × q*) matrix or data frame of read counts for *n* genes in *q* samples and `K=2:10` provides the desired range of numbers of clusters (here, 2 to 10). We note that this function directly calls the `Rmixmod` R package to fit Gaussian mixture models [32]. For backwards compatibility with our previous method [17], a similar function call may be used to fit a Poisson mixture model on raw counts using the `HTSCluster` package:

~~~
> run_pois <- coseq(counts, conds, K=2:10, model="Poisson")
~~~

where a vector `conds` is additionally provided to identify the experimental condition associated with each column in `counts`. In both cases, the output of the `coseq` function is an S4 object of class `coseqResults` (an extension of the `SummarizedExperiment0` Bioconductor S4 class) on which standard `plot` and `summary` functions can be directly applied; the former uses functionalities from the `ggplot2` package [37]. Several examples of the standard plot commands can be seen in the Results section of this work, as well as in the reproducible Rmarkdown document included in the Supplementary Materials. The option of parallelization via the `BiocParallel` Bioconductor package is also provided.

In addition to the choice of mixture model and transformation to be used, the `coseq` function provides flexibility to the user to filter normalized read counts according to their mean value if desired, specify library size normalization method (TMM, median ratio, upper quartile, or user-provided normalization factors), and modify `Rmixmod` options (number of iterations, etc). For the specific case of arcsin- and logit-transformed normalized profiles, we provide a convenience function `compareICL` to calculate and plot the corrected ICL model selection criteria defined in Equations (10) and (11). Finally, as RNA-seq expression analyses are often performed on a subset of genes identified as differentially expressed, the `coseq` function can also be directly called on an `DESeqResults` S4 object or integrated with `DGELRT` S4 objects, respectively corresponding to output from the `DESeq2` [11] and `edgeR` [10] Bioconductor packages for RNA-seq differential analyses. For more details and examples, see the full package vignette provided with `coseq`.

## Results

In the following, we illustrate co-expression analyses using Gaussian mixture models in conjunction with the proposed transformations on normalized expression profiles for several RNA-seq datasets. The data were selected to represent several different organisms (pig, mouse, human, fly) in studies for which co-expression is of particular interest (across tissues or across time); additional details on how data were obtained and preprocessed may be found in the Supplementary Materials.

### Description of RNA-seq data

#### Porcine small intestine

Mach et al. [38] used RNA-seq to study site-specific gene expression along the gastrointestinal tract of four healthy 70-day-old male Large White piglets. Samples were collected in three sites along the proximal-distal axis of the small intestine (duodendum, jejunum, and ileum), as well as the ileal Peyer’s patch (a lymphoid tissue localized in direct contact with the epithelial intestinal tissue). Complete information regarding sample preparation, sequencing, quality control, and pre-processing are available in the original article [38]. Raw reads are available at NCBI's SRA repository (PRJNA221286 BioProject; accessions SRR1006118 to SRR1006133); in the current work, read counts for genes sharing a common gene symbol or Ensembl gene ID were summed.

#### Embryonic mouse neocortex

Fietz et al. [29] studied the expansion of the neocortex in five embryonic (day 14.5) mice by analyzing the transcriptome of the ventricular zone (VZ), subventricular zone (SVZ), and cortical plate (CP) using RNA-seq. Laser-capture microdissection, RNA isolation and cDNA library preparation, and RNA sequencing and quantification are described in the Supplementary Materials of Fietz et al. [29]. In our work, raw read counts for this study were downloaded on December 23, 2015 from the Digital Expression Explorer (DEE) [39] using associated SRA accession number SRP013825, and run information was downloaded using the SRA Run Selector. Additional information about the DEE processing pipeline may be found in the Supplementary Materials.

#### Fetal human neocortex

In the aforementioned study, Fietz et al. [29] also included samples from 6 (13-16 wk postconception) human fetuses taken from four neocortex regions: CP, VZ, and inner and outer subventricular zone (ISZZ and OSVZ, respectively). Raw counts were obtained in the same manner as described above.

#### Dynamic expression in embryonic flies

As part of the modENCODE project to annotate functional elements of the *Drosophila melanogaster* genome, Graveley et al. [40] characterized the expression dynamics of the fly using RNA-seq over 27 distinct stages of development, from early embryo to ageing male and female adults. As in our previous co-expression work [17], we focus on a subset of these data from 12 embryonic samples that were collected at 2-hour intervals for 24 hours, with one biological replicate for each time point. Phenotype tables and raw read counts were obtained from the ReCount online resource [41].

### Results on RNA-seq data

We used the `coseq` package described in the previous section to fit Gaussian mixture models to the arcsin-and logit-transformed normalized profiles for each of the four datasets described above for *K* = 2*,…,* 40 clusters (with the exception of the *Drosophila melanogaster* data, for which a maximum value of *K* = 60 was used), using the TMM library size normalization, filtering genes with mean normalized count less than 50, and otherwise using default values for parameters. Concerning the filtering step, screening using either a differential analysis or a threshold on normalized means or coefficients of variation are often applied in practice prior to co-expression analyses to remove features that contribute noise. We note that for some studies (e.g., those in which some genes may be completely switched off in some conditions), a less stringent filtering threshold may be desired; in such cases, the `meanFilterCutoff` argument of `coseq` may be omitted (corresponding to no filter) or set to a smaller value. In all cases, we calculated the corrected ICL values from Equations (10) and (11) to compare between the arcsin and logit transformations; the number of clusters 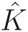 identified for each transformation, as well as the preferred model-transformation pair chosen via the corrected ICL, are shown for each dataset in Table 1. The corrected ICL values across a range of numbers of clusters *K* are shown in Figure 2 for the Graveley et al. [40] fly and Fietz et al. [29] mouse data; for clarity, we focus our discussion in the main text on these two datasets, but complete and reproducible results (in the form of an Rmarkdown document) for all four RNA-seq datasets may be found in the Supplementary Materials.

**Table 1:** Summary of results for Gaussian mixture models fit on transformed normalized RNA-seq profiles. For each dataset, the organism, associated reference, experimental conditions 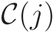 and number of biological replicates in each, and number of clusters selected via ICL for the arcsin and logit transformation are provided. Boldface values indicated the final model selected via the corrected ICL.

| Organism | Reference | Conditions (reps) | $\hat{K}_{\text{arcsin}}$ | $\hat{K}_{\text{logit}}$ |
| --- | --- | --- | --- | --- |
| Pig | Mach et al. [38] | 4 tissues (4 reps each) | **14** | 11 |
| Mouse | Fietz et al. [29] | 3 tissues (5 reps each) | **12** | 15 |
| Human | Fietz et al. [29] | 4 tissues (6 reps each) | **6** | 8 |
| Fly | Graveley et al. [40] | 12 time pts (1 rep each) | **28** | 23 |

**Figure 2:**
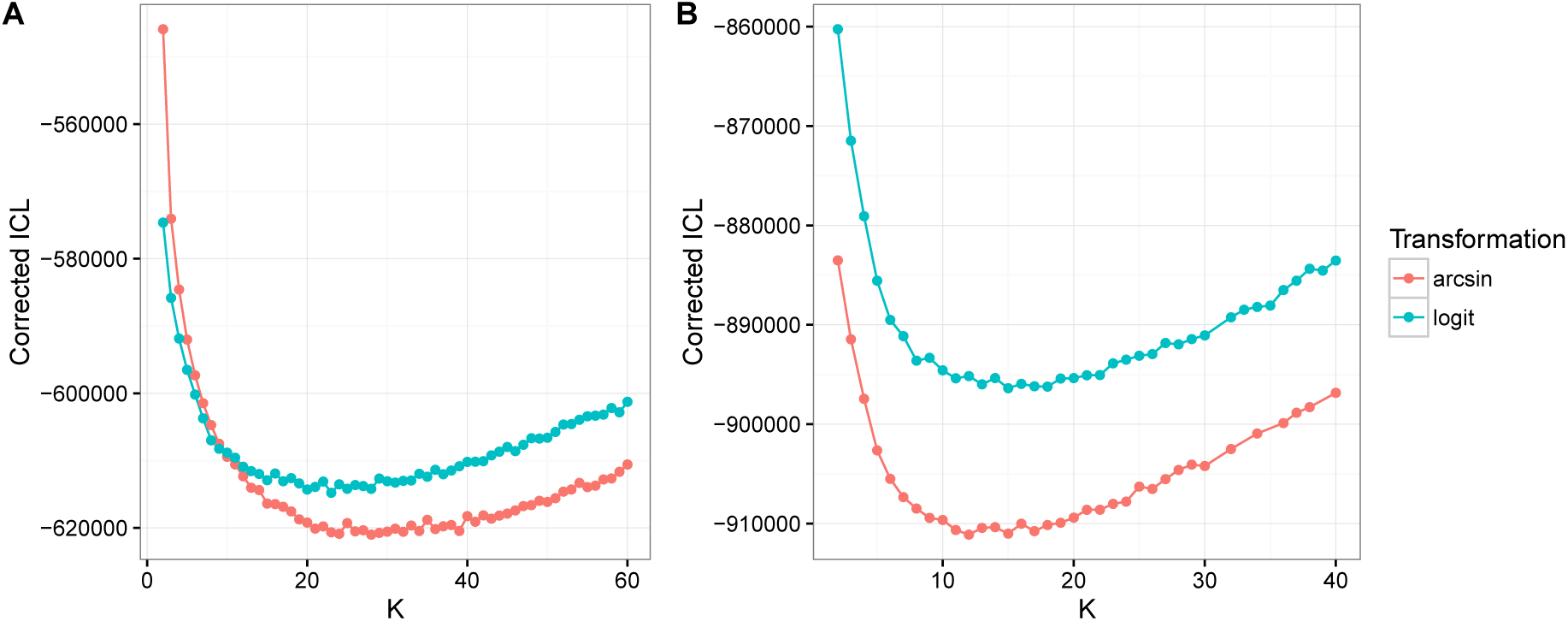
Corrected ICL values for the arcsin (red) and logit (blue) transformed normalized expression profiles over a range of numbers of clusters *K* for the Graveley et al. [40] fly and Fietz et al. [29] mouse data (A and B, respectively).

For the remainder of the article, the results presented correspond to the model selected via the corrected ICL. It is of interest to investigate the per-cluster covariance structures estimated for the selected models for each of the RNA-seq datasets. As an example, the per-cluster correlation matrices estimated by `coseq` for two selected clusters from the Graveley et al. [40] and Fietz et al. [29] mouse data are shown in Figure 3. It is interesting to note that although the Gaussian mixture model does not explicitly incorporate the experimental condition labels 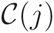, the estimated models include large cluster-specific correlations among close time points (Figures 3A and 3B) or among replicates within each tissue (Figures 3C and 3D). In addition, cluster-specific correlation structures among regions may be clearly seen; for example, in the Fietz et al. [29] mouse data, Cluster 2 is characterized by very large negative correlations between the CP and SVZ/VZ regions, while Cluster 3 instead has a strong negative correlation between the VZ and CP/SVZ regions. This strongly suggests that in these data, the assumption of conditional independence among samples assumed by the Poisson mixture model described in Rau et al. [17] is indeed unrealistic.

**Figure 3:**
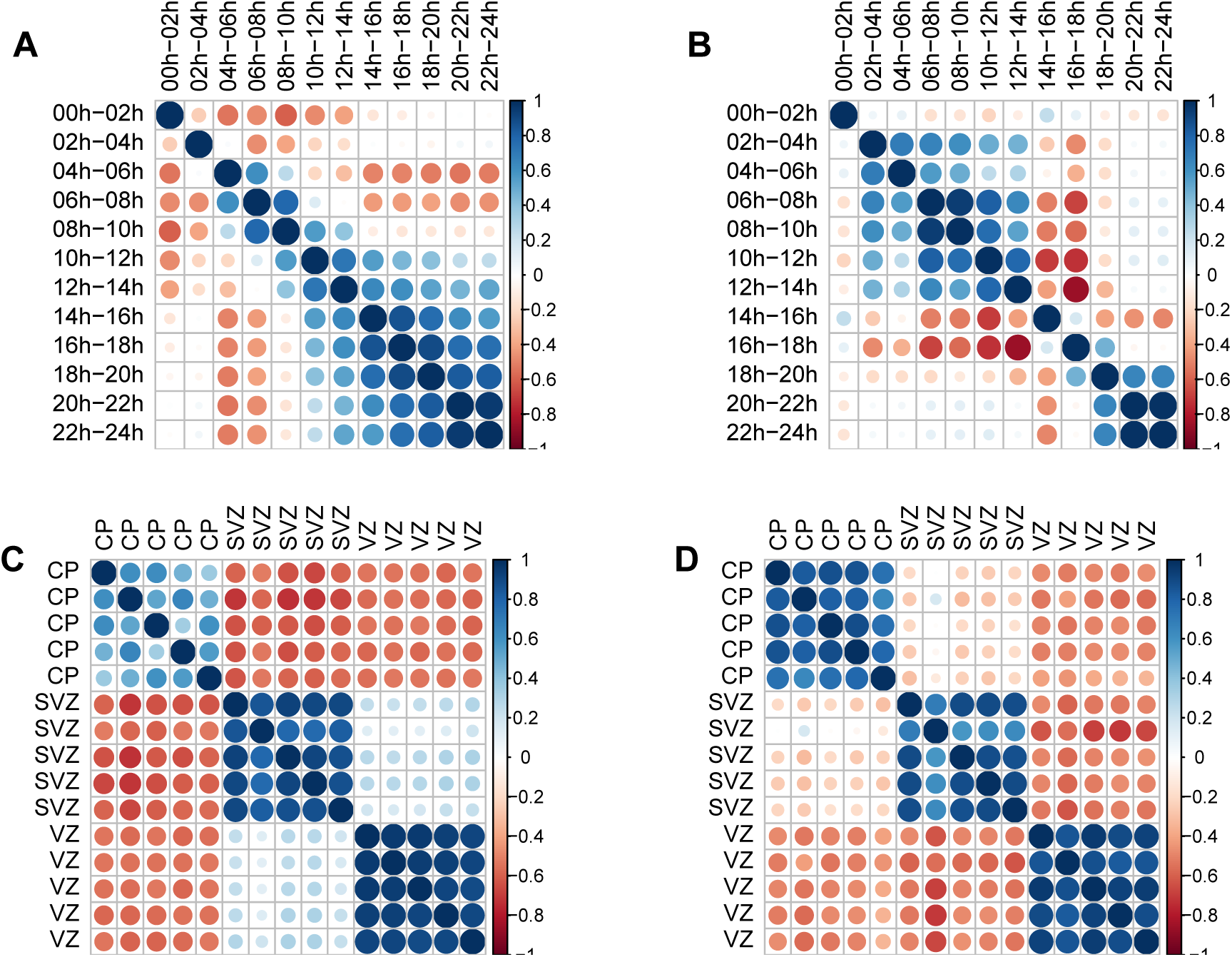
Per-cluster correlation matrices for clusters 25 (A) and 27 (B) from the Graveley et al. [40] fly data and for clusters 2 (C) and 3 (D) from the Fietz et al. [29] mouse data. Dark blue and red represent correlations close to 1 and −1, respectively, and circle areas correspond to the absolute value of correlation coefficients. Correlation matrices are visualized using the `corrplot` R package.

There are several ways in which per-cluster expression profiles can be represented graphically, depending on the type of data plotted (normalized counts, normalized expression profiles, or transformed normalized profiles), the type of plot (e.g., line plots or boxplots), and whether replicates within experimental conditions are averaged or plotted independently. Regarding the latter point, note that the Gaussian mixture model is fit on the entirety of the data, and replicate averaging is proposed to simplify the visualization of cluster-specific expression. Although the `coseq` package facilitates the implementation of any combination of these three graphical options, our recommendations for visualizating co-expression results are as follows: 1) although “tighter" profiles are observed when plotting the transformed normalized profiles (as these are the data used to fit the model), interpretation of profiles is improved by instead using the untransformed normalized profiles; 2) boxplots are generally preferable when experimental conditions 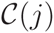 represent distinct groups, although line plots can be useful for time-course experiments; 3) averaging replicates prior to plotting often provides clearer distinctions among cluster-specific profiles. Following these recommendations, the cluster-specific profiles identified for the Graveley et al. [40] and Fietz et al. [29] mouse data are shown in Figures 4 and 5.

**Figure 4:**
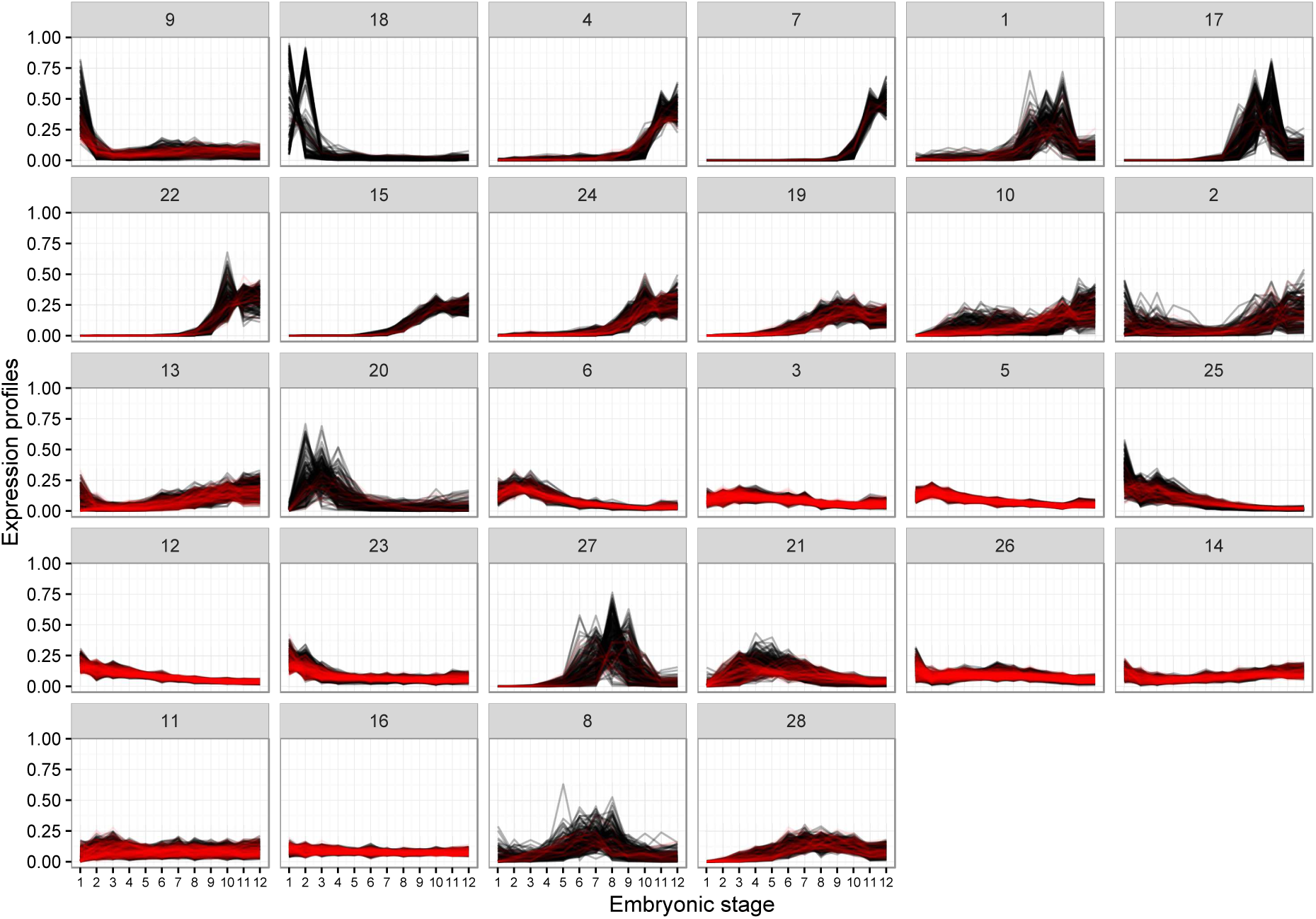
Per-cluster expression profiles for the Graveley et al. [40] data. Clusters have been sorted so that those with similar mean vectors (as measured by the Euclidean distance) are plotted next to one another. Red lines correspond to genes with maximimum conditional probability *τ*_max_(*i*) of cluster membership < 0.8.

**Figure 5:**
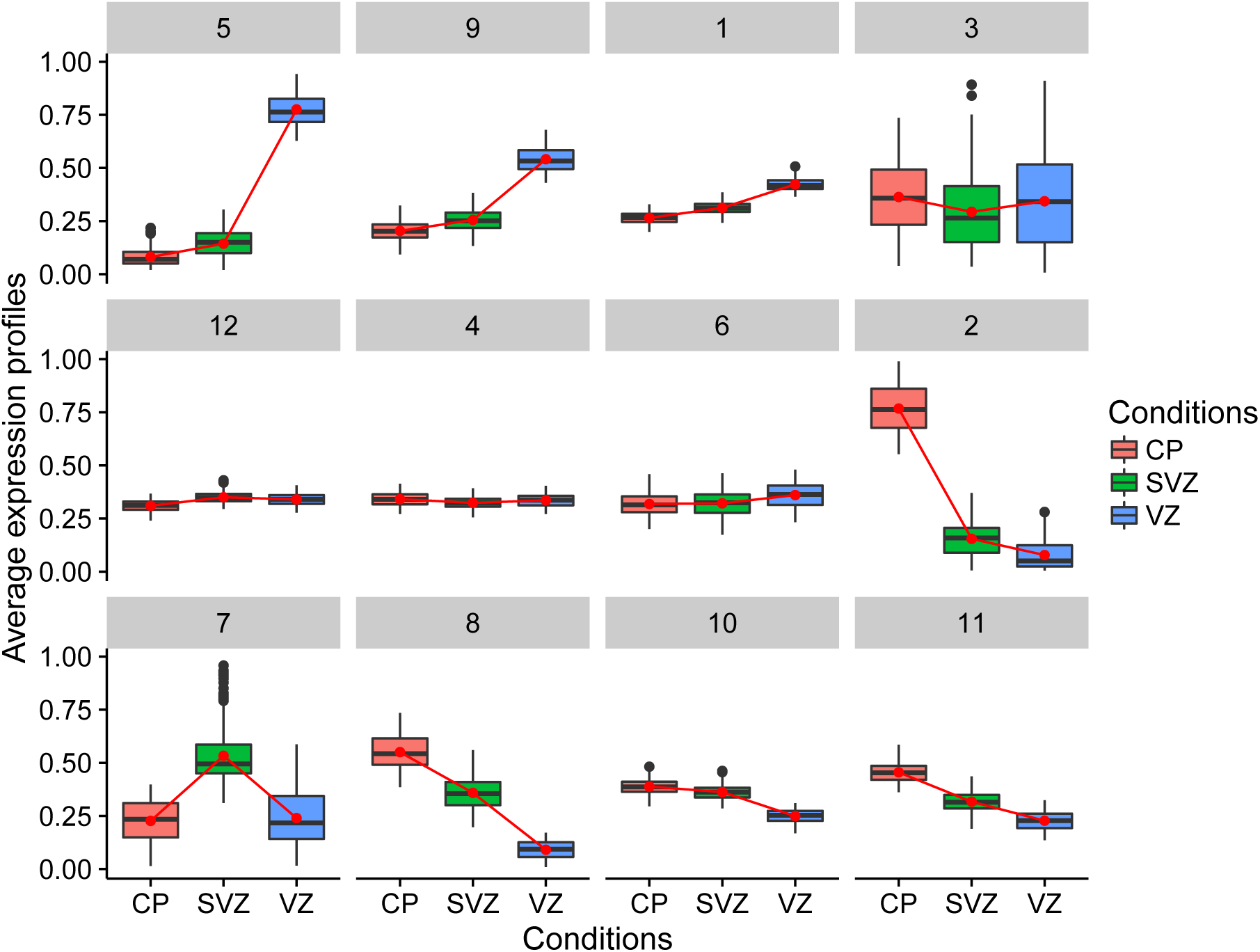
Per-cluster expression profiles for the Fietz et al. [29] data. Clusters have been sorted so that those with similar mean vectors (as measured by the Euclidean distance) are plotted next to one another. Connected red lines correspond to the mean expression profile for each group.

An additional advantage of model-based clustering approaches is that they facilitate an evaluation of the clustering quality of the selected model by examining the maximum conditional probabilities of cluster membership for each gene:

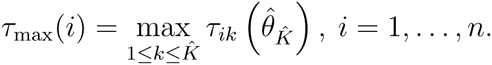

Boxplots of the maximum conditional probabilities *τ*_max_(*i*) per cluster for the Graveley et al. [40] and Fietz et al. [29] mouse data are presented in Figure 6. It may be seen that across clusters, the majority of genes in both datasets have a large value (i.e., close to 1) for *τ*_max_(*i*); the number of genes with *τ*_max_(*i*) > 0.8 is 7822 (82.1%) and 7382 (82.4%) for the Graveley et al. [40] and Fietz et al. [29] mouse data, respectively. However, the boxplots also illustrate that some genes have a *τ*_max_(*i*) less than this threshold, in some cases as low as 0.4; this indicates that for a small number of genes, the cluster assignment is fairly ambiguous and assignment to a single cluster is questionable (the gene with the smallest *τ*_max_(*i*) in the Fietz et al. [29] mouse data had a conditional probability of 24.8%, 32.2%, 13.0% and 30.0% of belonging to clusters 1, 4, 6, and 12, respectively). In such cases, it may be prudent to focus attention on genes with highly confident cluster assignments (e.g., those with *τ*_max_(*i*) > 0.8).

**Figure 6:**
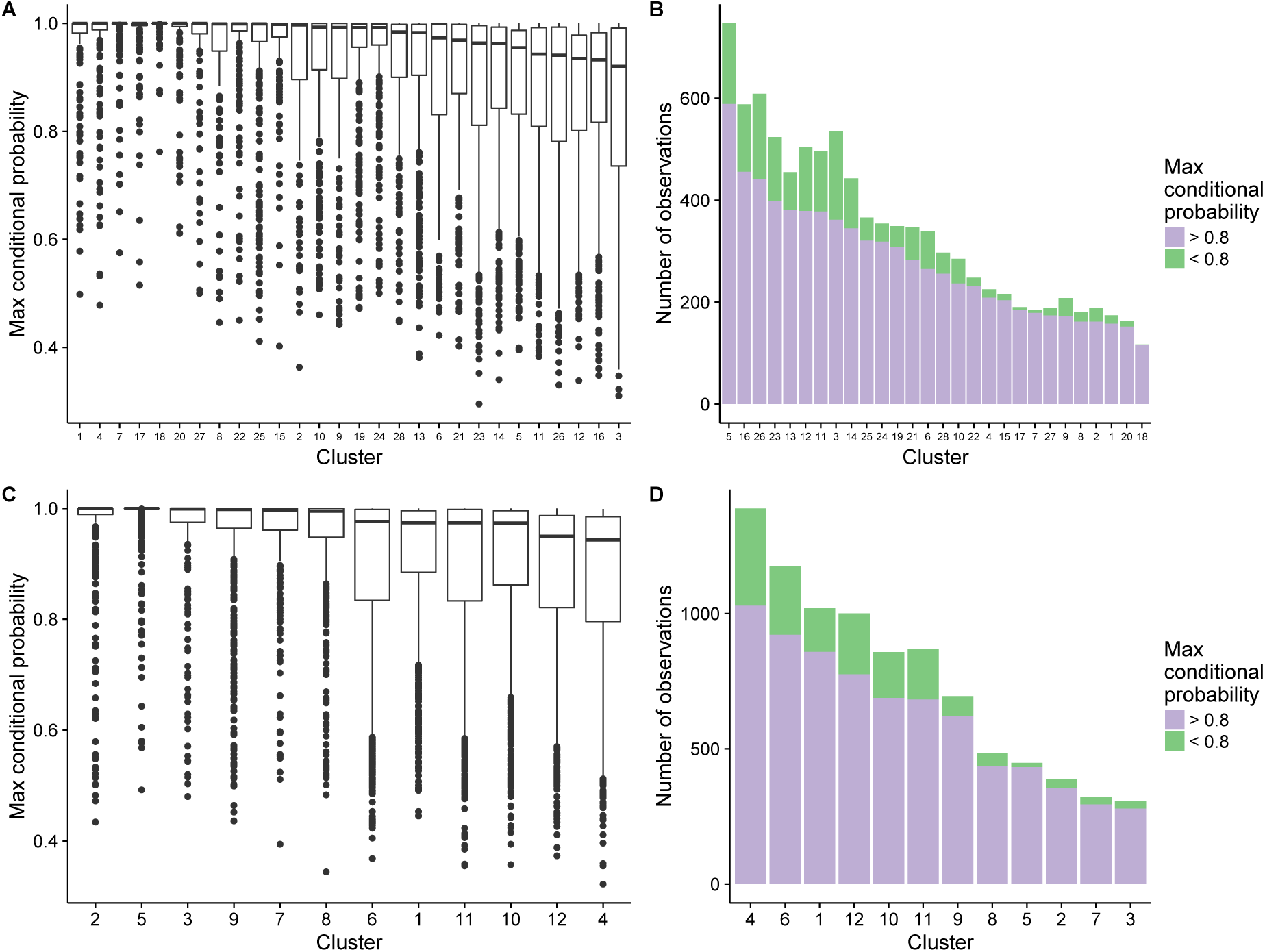
Evaluation of clustering quality for the Graveley et al. [40] (top) and Fietz et al. [29] mouse (bottom) data. (left) Maximum conditional probabilities *τ*_max_(*i*) for each cluster, sorted in decreasing order by the cluster median. (right) Barplots of cluster sizes, according to *τ*_max_(*i*) greater than or less than 0.8, sorted according to the number of genes with *τ*_max_(*i*) > 0.8.

Finally, examining the distribution of *τ*_max_(*i*) values within each cluster provides information about the homogeneity and relevance of each cluster. For both datasets, all cases clusters are primarily made up of genes with highly confident *τ*_max_(*i*) values; however some clusters (e.g., Clusters 1 and 4 in the Graveley et al. [40] data) appear to be more homogeneous and well-formed than others (e.g., Clusters 16 and 3 in the same data). These conclusions align with the general observations made about the per-cluster normalized profiles in Figure 4, where it may be seen that clusters 16 and 3 have quite similar profiles (suggesting that unambiguous assignment to one of these clusters is more difficult).

## Discussion and recommendations

In this work, we have primarily addressed the choice of data to be clustered (transformed normalized profiles rather than raw counts) for RNA-seq co-expression analysis under the framework of Gaussian mixture models. In the following section, we provide some additional discussion of the choice of transformation and the use of Gaussian mixture models for co-expression analysis, as well as additional remarks about the practical application of co-expression analyses.

### Choice of transformation for co-expression analysis

We have focused the majority of our discussion here on the use of (arcsin- or logit-transformed) normalized profiles to identify groups of co-expressed genes. As previously noted, a variety of well-established candidates for transformations have already been proposed for RNA-seq data, including the log(· + *c*) (for a constant *c*), VST, CPM, and rlog. As suggested by a reviewer, we could consider the alternative approach of clustering RNA-seq data after applying one of these transformations and mean-centering per gene; this latter step would remove the spread in values observed, for example, in the log-transformed (and uncentered) data in Figure 1B and ensure that the data to be clustered are independent of the absolute expression levels.

To investigate this idea, we fit Gaussian mixture models to the centered log, VST, CPM, and rlog transformed data for the Fietz et al. [29] mouse data with *K* = 2, …, 40 clusters using the same parameters as for the `coseq` analysis. For all four mean-centered transformations across all clusters, the most general covariance model form (the so-called [*p*_*k*_*L*_*k*_*C*_*k*_] model, with variable proportions, volume, orientation, and shape) systematically resulted in estimation errors via the Rmixmod package. When considering a larger variety of general covariance model forms (the [*p*_*k*_*LC*_*k*_], [*p_k_L_k_C*], [*p_k_LC*] forms, which all have variable proportions and respectively feature equal volume, equal orientation/shape, and equal volume/orientation/shape), the same estimation errors were observed for all centered transformations across all clusters. We were in fact only able to estimate the Gaussian mixture model for all four centered transformations when restricting the covariance model forms to be spherical or diagonal (which roughly corresponds to applying a K-means type algorithm); as such, this implies that when using these alternative transformations, complex covariance structures among samples (such as those observed in Figure 3) must be assumed to be negligeable (i.e., conditional independence given the clustering component).

### Gaussian mixture models for co-expression analysis

We have illustrated several advantages in using Gaussian mixture models in conjunction with appropriately defined transformations to identify groups of co-expressed genes from normalized RNA-seq gene expression profiles. First, mixture models in general have the advantage of providing a rigorous statistical framework for parameter estimation, an objective assessment of the number of clusters present in the data through the use of penalized criteria, and the possibility of performing diagnostic checks on the quality and homogeneity of the resulting clusters. In particular, diagnostic plots on the maximum conditional probabilities of cluster membership provide a global overview of the clustering and an objective explanation of the quality of cluster assignments. Since only a subset of genes are expected to be assigned to biologically interpretable groups, these diagnostic plots help provide a basis for discussion about the choice of genes for follow-up study. However, it should be noted that such assessments of cluster stability are not unique to Gaussian mixture models. In recent years, several resampling-based methods for assessing cluster stability and membership have been proposed, including the `clue` R package [42] and `ConsensusClusterPlus` Bioconductor package [43]; for example, Ohnishi et al. [44] applied K-means clustering to resampled microarray data, constructed a consensus clustering using `clue`, and compared each of the resampled clusterings to the consensus to identify stable and robust clusters.

Gaussian mixtures in particular represent a rich, flexible, and well-characterized class of models that have been successfully implemented in a large variety of theoretical and applied research contexts. For RNA-seq data, this means that the model may directly account for per-cluster correlation structures among samples, which can be quite strong in RNA-seq data. In this work we considered a single form of Gaussian covariance matrices (the [*p*_*k*_*L*_*k*_*C*_*k*_] form), but any or all of the 28 forms of Gaussian mixture models could be used in practice.

We have also discussed the use of penalized criteria like the ICL and BIC to objectively compare results between different transformations, and potentially among different forms of Gaussian covariance matrices or among different models. For the four datasets considered here, the arcsin transformation of normalized expression profiles was consistently preferred to the logit transformation; as previously mentioned, this is likely due the sensitivity of the latter to very small *p*_*ij*_ values. An interesting further direction of research would be to consider approaches able to directly model the compositional nature of normalized profiles *p*_*ij*_ without the need to apply an arcsin or logit transformation.

### Practical issues for co-expression analysis

It is worth noting that the concept of *gene co-expression* is alternatively used to refer to two broad types of analyses [45]: 1) clustering gene expression patterns to explore shared function and co-regulation (our focus in this work); and 2) network inference, which aims to construct a model of the network of regulatory interactions between genes. For the latter, a popular and widely-used method is the weighted correlation network analysis (WGCNA) approach [46], which seeks to identify modules of highly interconnected (both positively and negatively correlated) genes. Although this approach was first proposed for use with microarray data, the WGCNA online FAQ page suggests it may be used for normalized RNA-seq data following a variance-stabilizing or log transformation; however, as WGCNA is based on esimates of pair-wise correlation among genes, the authors recommend at least 15 to 20 samples in practice (as a reminder, the number of samples in the four datasets considered here ranged from 12 to 24). As such, due to both the difference in analysis objectives (clustering versus network inference) and the relatively small sample sizes of the four datasets, we have not included a more detailed discussion of WGCNA in the current work.

Many alternative clustering strategies exist based on different algorithms (e.g., K-means and hierarchical clustering), distance measures calculated among pairs of genes (e.g., Euclidean distance, correlation, etc), and techniques for identifying the number of clusters (e.g., the Dynamic Tree Cut method for dendrograms [46]). The difficulty of comparing clusterings arising from different approaches is well-known, and it is rarely straightforward to establish the circumstances under which a given strategy may be preferred. One possibility that may be of interest in practice is to analyze cluster ensembles arising from a set of different methods to assess the agreement or dissimilarity among partitions and obtain a consensus clustering [42].

In addition to the choice of clustering method, several practical issues should be considered in co-expression analyses. First, a common question is whether genes should be screened prior to the analysis (e.g., via an upstream differential analysis or filter based on the mean expression or coefficient of variation for each gene). Such a screening step is often used in practice, as genes contributing noise but little biological signal of interest can adversely affect clustering results. A second common question pertains to whether replicates within a given experimental group should be modeled independently or summed or averaged prior to the co-expression analysis. Although technical replicates in RNA-seq data are typically summed prior to analysis, in this work we fit Gaussian mixture models on the full data including all biological replicates; subsequently to visualize clustering results, replicate profiles are averaged for improved clarity of cluster profiles.

Following a co-expression analysis, it is notoriously difficult to validate the results of a clustering algorithm on transcriptomic data, and such results can be evaluated based on either statistical criteria (e.g., between-group and within-cluster inertia measures) or external biological criteria. In practice groups of co-expressed genes are further characterized by analyzing and integrating various resources, such as functional annotation or pathway membership information from databases like the Gene Ontology Consortium. Such functional analyses can be useful for providing interpretation and context for the identified clusters.

## Key Points

- After applying an appropriate transformation, Gaussian mixture models represent a rich, flexible, and well-characterized class of models to identify groups of co-expressed genes from RNA-seq data. In particular, they directly account for per-cluster correlation structures among samples, which are observed to be quite strong in typical RNA-seq data.
- Normalized expression profiles, rather than raw counts, are recommended for co-expression analyses of RNA-seq data. Because these data are compositional in nature, an additional transformation (e.g., arcsin or logit) is required prior to fitting a Gaussian mixture model.
- Penalized model selection criteria like the BIC or ICL can be used to select both the number of clusters present and the appropriate transformation to use; in the latter case, an additional term based on the Jacobian of the transformation is added to the criterion, yielding a corrected BIC or ICL that can be used to directly compare two transformations.

## Acknowledgements

The authors would like to thank Gilles Celeux, Sandrine Laguerre, Béatrice Laurent, Clément Marteau, and, Marie-Laure Martin-Magniette for helpful discussions. We also thank the three reviewers for their constructive comments that helped to improve this work.

## References

[1] M. B. Eisen et al. Cluster analysis and display of genome-wide expression patterns. PNAS, 95(25):14863–14868, 1998.

[2] D. Jiang, C. Tang, and A. Zhang. Cluster analysis for gene expression data: A survey. IEEE Transactions on Knowledge and Data Engineering, 16(11):1370–1386, 2004.

[3] K. Y. Yeung et al. Model-based clustering and data transformations for gene expression data. Bioinformatics, 17(10):977–987, 2001.

[4] S. Anders, P.T. Pyl, and W. Huber. A Python framework to work with high-throughput sequencing data. Bioinformatics, 31(2):166–169, 2015.

[5] Y. Liao, G. K. Smyth, and W. Shi. featureCounts: an efficient general purpose program for assigning sequence reads to genomic features. Bioinformatics, 30(7):923–930, 2014.

[6] B. Li and C. N. Dewey. RSEM: accurate transcript quantification from RNA-Seq data with or without a reference genome. BMC Bioinformatics, 12(323), 2011.

[7] N. L. Bray et al. Near-optimal probabilistic RNA-seq quantification. Nature Biotechnology, 34:525–527, 2016.

[8] A. Oshlack and M. J. Wakefield. Transcript length bias in RNA-seq data confounds systems biology. Biology Direct, 4(14), 2009.

[9] P. P. Łabaj et al. Characterization and improvement of RNA-Seq precision in quantitative transcript expression profiling. Bioinformatics, 27(ISMB):i383–i391, 2011.

[10] M. D. Robinson et al. edgeR: a Bioconductor package for differential expression analysis of digital gene expression data. Bioinformatics, 26:139–140, 2010.

[11] M. I. Love, W. Huber, and S. Anders. Moderated estimation of fold change and dispersion for RNA-seq data with DESeq2. Genome Biology, 15(550), 2014.

[12] J. H. Ward. Hierarchical grouping to optimize an objective function. Journal of the American Statistical Association, 58(301):236–244, 1963.

[13] J. B. MacQueen. Some methods for classification and analysis of multivariate observations. In Proceedings of the 5th Berkeley Symposium on Mathematical Statistics and Probability, number 1, pages 281–297. Berkeley, University of California Press, 1967.

[14] G. McLachlan and D. Peel. Finite Mixture Models. Wiley-Interscience, 2000.

[15] D. M. Witten. Classification and clustering of sequencing data using a Poisson model. Annals of Applied Statistics, 5(4):2493–2518, 2011.

[16] S. Anders and W. Huber. Differential expression analysis for sequence count data. Genome Biology, 11(R106):1–28, 2010.

[17] A. Rau et al. Co-expression analysis of high-throughput transcriptome sequencing data with poisson mixture models. Bioinformatics, 31:1420–1427, 2015.

[18] G. Schwarz. Estimating the dimension of a model. The Annals of Statistics, 6(2):461–464, 1978.

[19] C. Biernacki, G. Celeux, and G. Govaert. Assessing a mixture model for clustering with the integrated completed likelihood. IEEE Transactions on Pattern Analysis and Machine Intelligence, 22(7):719–725, 2000.

[20] L. Birgé and P. Massart. Gaussian model selection. J. Eur. Math. Soc., 3:203–268, 2001.

[21] J.-P. Baudry et al. Slope heuristics: overview and implementation. Stat. Comp., 22:455–470, 2012.

[22] Y. Si et al. Model-based clustering for RNA-seq data. Bioinformatics, 30(2):197–205, 2014.

[23] C.W. Law et al. voom: precision weights unlock linear model analysis tools for RNA-seq read counts. Genome Biology, 15(R29), 2014.

[24] G. K. Smyth. Linear models and empirical Bayes methods for assessing differential expression in microarray experiments. Statistical Applications in Genetics and Molecular Biology, 1(3):1–26, 2004.

[25] M. D. Robinson and A. Oshlack. A scaling normalization method for differential expression analysis of RNA-seq data. Genome Biology, 11(R25), 2010.

[26] M.-A. Dillies et al. A comprehensive evaluation of normalization methods for Illumina high-throughput RNA sequencing data analysis. Briefings in Bioinformatics, 14(6):671–683, 2013.

[27] R. Tibshirani. Estimating transformations for regression via additivity and variance stabilization. Journal of the American Statistical Association, 83:394–405, 1988.

[28] W. Huber et al. Parameter estimation for the calibration and variance stabilization of microarray data. Statistical Applications in Genetics and Molecular Biology, 2(1):Article 3, 2003.

[29] S. A. Fietz et al. Transcriptomes of germinal zones of human and mouse fetal neocortex suggest a role of extracellular matrix in progenitor self-renewal. PNAS, 109(29):11836–11841, 2012.

[30] J. Aitchison. The Statistical Analysis of Compositional Data. Chapman & Hall, 1986.

[31] G. Celeux and G. Govaert. Gaussian parsimonious clustering models. Pattern Recognition, 28(5):781 – 793, 1995.

[32] R. Lebret et al. Rmixmod: The R Package of the model-based unsupervised, supervised, and semi-supervised classification Mixmod library. Journal of Statistical Software, 67(6): 1–29, 2015.

[33] A. P. Dempster et al. Maximum likelihood from incomplete data via the EM algorithm. Journal of the Royal Statistical Society, Series B (Methodological), 39(1):1–38, 1977.

[34] L. Birgé and P. Massart. Minimal penalties for Gaussian model selection. Probability Theory and Related Fields, 138:33–73, 2007.

[35] I. Thomas, P. Frankhauser, and C. Biernacki. The fractal morphology of the built-up landscape. Landscape of Urban Plan, 84(2):99–115, 2008.

[36] M. Gallopin. Classification et inférence de réseaux pour les données RNA-seq. PhD thesis, Université Paris-Saclay, 2015.

[37] Hadley Wickham. ggplot2: Elegant Graphics for Data Analysis. Springer-Verlag New York, 2009. URL http://ggplot2.org.

[38] N. Mach et al. Extensive expression differences along porcine small intestine evidenced by transcriptome sequencing. PLoS ONE, 9(2):1–12, 02 2014.

[39] M. Ziemann et al. Digital Expression Explorer: A user-friendly repository of uniformly processed RNA-seq data. In ComBio2015, volume POS-TUE-099, Melbourne, 2015. doi: 10.13140/RG.2.1.1707.5926.

[40] B. R. Graveley et al. The development transcriptome of *Drosophila melanogaster*. Nature, 471:473–479, 2011.

[41] A. C. Frazee, B. Langmead, and J. T. Leek. ReCount: a multi-experiment resource of analysis-ready RNA-seq gene count datasets. BMC Bioinformatics, 12, 2011.

[42] K. Hornik. A CLUE for CLUster Ensembles. Journal of Statistical Software, 14(12), 2005.

[43] M. D. Wilkerson and D. N. Hayes. ConsensusClusterPlus: a class discvoery tool with confidence assessments and item tracking. Bioinformatics, 26(12):1572–1573, 2010.

[44] Y. Ohnishi et al. Cell-to-cell expression variability followed by signal reinforcement progressively segregates early mouse lineages. Nature Cell Biology, 16:27–37, 2014.

[45] P. D’haeseleer, S. Liang, and R. Somogyi. Genetic network inference: from co-expression clustering to reverse engineering. Bioinformatics, 16(8):707–726, 2000.

[46] P. Langfelder and S. Horvath. WGCNA: an R package for weighted correlation network analysis. BMC Bioinformatics, 9(559), 2008.

